# Impact of Regularization Methods and Outlier Removal on Unsupervised Sample Classification

**DOI:** 10.64898/2026.04.07.716815

**Authors:** Carol Heckman

## Abstract

**Background:** High-content assays (HCAs) have problems distinguishing biologically significant effects from the incidental effects of non-repeatable technical factors. Non-repeatable results are attributed to variations in the cell culture environment and the numerous, heterogeneous descriptors evaluated. The aim here was to determine whether preprocessing operations impacted the reproducibility of class assignments of experimental data.

**Methods:** Batch effects that could affect reproducibility, i.e., signal/noise ratio, instrumental conditions, and segmentation, were controlled variables. The remaining batch effects, variations in materials, personnel, and culture environment could not be controlled. Descriptors’ values were measured directly from images. Exploratory factor analysis was used to solve the identifiable and interpretable feature, factor 4. In each of five trials, one sample was treated with the same chemical mixture (EXP) and another with the solvent vehicle alone (CON).

**Results:** Repeated CON and EXP samples showed significant differences among factor 4 means in data regularized within each trial. The mean of Trial 3 CON differed significantly from all other CON samples. These differences disappeared upon regularization to comprehensive databases. Among repeated EXPs, the Trial 2 mean differed from three other EXPs, but regularization to comprehensive databases had little effect. However, classification patterns were unchanged after regularization to any comprehensive database derived by the same protocol. After regularization to datasets derived by two different protocols, the classification pattern differed but only reflected elevation of differences that had been marginal to statistical significance. Outlier removal was deleterious. Even with the most sparing definition of outliers, over 3% of a single sample’s contents were removed from most trials. Elimination based on the overall within-trial distributions caused type I and type II errors.

**Conclusions:** Non-repeatable factor 4 means in repeated trials had negligible influence on classification outcomes, so repeatability may not be a good indicator of assay quality. Irreducible batch effects, combined with small sample sizes and skewed distributions of descriptors’ values, may account for non-repeatability. As the current results are based on real-world data, they suggest that non-repeatability is an uncorrectable feature of these assays. Classification patterns are not affected by several irreducible technical factors, namely materials, personnel, and non-repeatable environmental variables.

## 1 Introduction

Methods of interpreting images are under investigation in many scientific fields, but they are particularly critical in cell biology where much of the information used to develop and test a hypothesis depends on images. A common concern of scientists in many fields is how to ensure that information extracted from images is correctly interpreted. One solution is to use “supervised learning” procedures in which machine models are trained by human arbiters to recognize objects of interest. The power of this approach was demonstrated by the development of an automated process for detecting abnormal cells in cervical smears. Fourier descriptors were combined with image-based geometrical descriptors and textural measures [1]. Then, their values were sent through a convolutional neural network based on multilayer perceptron concepts to solve the pattern recognition problem [2, 3], (see for review [4]). However, the machine learning procedure only rivaled the performance of a human expert if the processes of sample collection and staining were also standardized [5].

Although the supervised methods allow one to extract valuable details from the sample or recognize objects with rare characteristics, human arbiters may have different standards for recognition of the imaged objects. In addition to bias, another drawback is the time and effort that humans must expend to analyze or classify large image datasets. Perhaps the main drawback of supervised learning, however, is that some important features may not be known to a human observer. These methods are particularly ill-suited for projects where workers want to recognize phenotypes relevant to drug development but lack a detailed knowledge of the mechanisms causing the disease state [6–9]. Instead, investigators acquire large numbers of images and assemble the outputs into multivariate datasets for “unsupervised” learning, in which all steps of selection, imaging, and processing are algorithmic. Complex, multivariate datasets of this type must subsequently be analyzed by statistical methods.

Measuring the values of descriptors directly from the image is a common starting point. Their number could range from a few dozen [10–12] to hundreds [6, 13] or even thousands [14, 15], all representing some aspect of the sample. Descriptors may measure complementary, redundant, or even contradictory aspects of the object, so, in a large dataset, much of the information is correlated. To reduce the number, descriptors with better predictive ability are selected for use, or they are combined into a new variable via mathematical manipulations. The latter methods avoid eliminating information; rather, they project the same information onto fewer variables. While it is widely held that low-dimensional variables, such as principal components and latent factors, are less interpretable than the descriptors from which they are derived [16, 17], others suggest that factors are more interpretable [10, 18–20]. This may be due to the tendency of factor rotation to perform statistical inference by orthogonalization of the multidimensional space [21].

This laboratory has placed considerable reliance on exploratory factor analysis to identify the properties and traits of cells prepared by label-free techniques. When cell protrusions were analyzed, based on orthogonalized descriptors, some of the low-dimensional variables showed a one-to-one correspondence to well-known protrusions of cells in culture [19, 22–24]. The current study employs one such latent factor as the trait or property, factor 4, so-called because it accounted for the fourth greatest fraction of the total variability in experimental datasets. It represents the filopodium, a protrusion type recognized by algorithms created independently in several laboratories [25–27]. The similarity among the laboratory’s definitions suggests it is a canonically isomorphic, or identifiable, object. The filopodia measurements rise and fall according to physiological conditions that augment or inhibit the structure’s prevalence in cultured cells [28–31], which suggests that it is also interpretable. As this implies reproducibility, such as variable is ideal for investigating the effect of procedures for processing unsupervised data workflows. To this end, data from five experiments were compared with respect to filopodia. In each experiment, there was one group of cells treated for 10 hours with 12-myristate 13-acetate (PMA) and 1-oleoyl-sn-glycero-3-phosphate (lysophosphatidic acid, LPA) and another treated with the solvent vehicle alone. Each sample consisted of 27-37 single cells in culture, and each cell’s image was reduced to a silhouette before being subjected to processing (see Materials and Methods). Latent factors were derived as illustrated in Figure 1.

**Figure 1.**
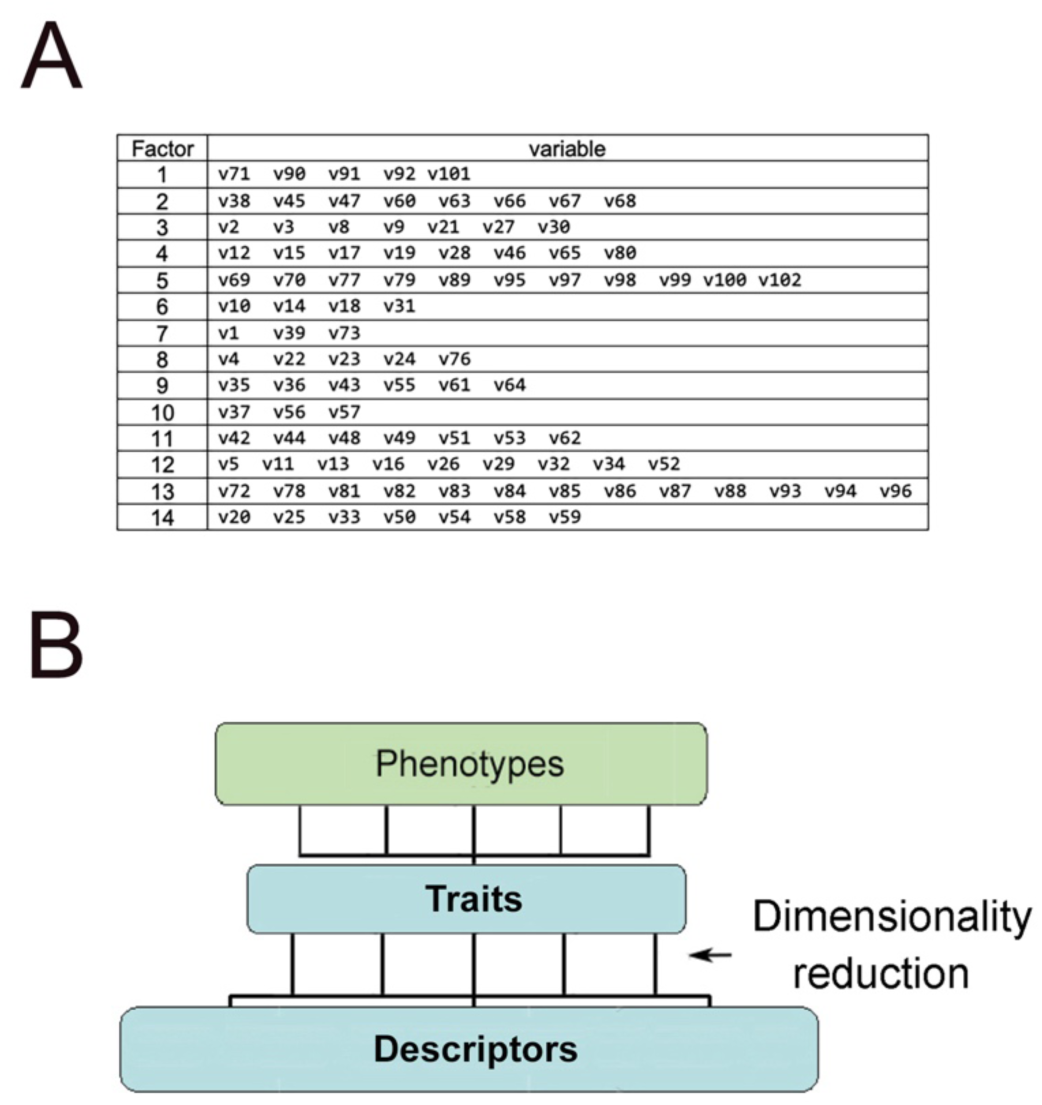
Relationship among descriptors, traits, and phenotypes. (A) The values of descriptors, called features by some earlier workers, are calculated based on attributes of each image. Actual cell features like the protrusions, filopodia, lamellipodia, and neurites, are called traits. The traits are derived from the correlation matrix of the descriptors’ values using latent factor analysis [44]. Here, the reproducibility of experiments is tested using factor 4. (B) The traits represent identifiable structural aspects of a cell. Traits are combined in different patterns to make up phenotypes (see 4.4 Classification).

The research aimed to determine how a class assignment of real-world experimental data could be made repeatable. The data were well-suited to addressing reproducibility because the assay protocol allowed many of the technical problems to be minimized. Batch effects or poor image quality may be hard to detect or compensate [32, 33], but can lead to irreproducible results. Here, the origin of the silhouettes depended on only one physical basis, and the boundaries were extracted by exact segmentation. This minimized variations in images due to instrument settings and eliminated concerns about signal/noise ratio. The problem of batch effects, i.e., non-repeatable results of a single treatment under ostensibly similar conditions (see for review [34, 35]), is reduced in this study to differences among the means of similar samples from repeated trials at a P-value <0.05 (see 2.1 Data generation). The approach was described in a previous report [36].

## 2 Materials and Methods

### 2.1 Data generation

Five sequential trials were run in the same laboratory, but at different times and by different personnel using different lots of reagents. The panel of drugs tested was not exactly reproduced, but one sample was treated with solvent vehicle alone, and another was treated for 10 hours with phorbol 13-myristate 12-acetate (PMA) and lysophosphatidic acid (LPA). A sample consisted of silhouettes of 27-36 cells of a treated or control group, and Trials 1-5 comprised a total of 46 samples. The steps of data acquisition, as described previously [19], were: 1) acquire images of single cells by scanning electron microscopy, 2) trace the boundary, and 3) compute a total of 33 dimensionless descriptors’ values. A factor 4 value for each cell was computed from a linear combination of descriptors’ weighted values, as shown in Figure 1. Factor extraction finds latent structures that are not directly measurable from the images but explain patterns observed in the data.

Under these conditions, PMA and LPA had no effect on factor 4 values compared to the control. Trial 1 (CON1 and EXP1) was described in [19], Trial 2 in [23], and the remaining samples in [22]. The differences between CON and EXP samples were not addressed in previous reports unless pertinent to the findings. Therefore, a preliminary analysis was undertaken to confirm that PMA- and LPA-treated samples resembled the controls. The ten samples were compared using the Kruskal-Wallis test with Bonferroni correction. Results of post-hoc Dunn’s test for differences among groups were obtained at <htpps://www.statskingdom.com>. As shown in Supplementary Material (Figure S1), there were 13 differences in total, but the control and treated samples never differed significantly within a single trial.

Because the CON samples were statistically indistinguishable from those treated with PMA and LPA, the experiments were ideal for studying the origin of false negative and false positive results. Specifically, the relationship between preprocessing and the reproducibility of class assignments could be investigated. The scoring coefficients used to calculate the factor values were derived from a database of 800 cells that varied in tissue origins and differentiated states. These data were mean-centered and scaled by the standard deviation before computing the coefficients [37]. Mean-centering adjusted the mean of the total values to zero. Scaling was done by dividing these values by the standard deviation of all values in the dataset. This process, known as autoscaling, normalization, regularization, StandardScaler, or z-scoring, is typically incorporated into workflows for handling microscopic images. Data from Trials 1-5 were generated by hand, i.e. exact tracings of cell boundaries from images taken at high magnification. As this ensured accurate quantification of the trait, the issue of classifying each sample relative to the others could be investigated without undue concern about batch effects arising from the procedures. The simplification of the protocol allowed the investigator to deal with questions of regularization and data cleaning as well as the statistical errors that could interfere with repeatability.

### 2.2 Data evaluation

To visualize the distribution of factor 4 in the paired control (CON) and treated (EXP) samples, values were plotted using the online site, <https://huygens.science.uva.nl/PlotsOfData/>. Whether differences between the factor 4 means of samples within a trial reached P<0.05 was determined by one-way analysis of variance (ANOVA), using the online service <https://www.statskingdom.com> with the recommended defaults. The settings, which can be reproduced in other statistical analysis software, are shown in Supplementary Material (Figure S2A). The rank-based non-parametric Kruskal-Wallis test was performed as described above to determine whether there were any differences among the repeated samples of Trials 1-5. The settings for the test are shown in Supplementary Material (Figure S2B).

Outliers were identified by calculating the interquartile range, IQR=Q3−Q1, for each sample. The values, 0.25–1.5 × IQR and 0.75+1.5 × IQR, were set as the lower and upper bounds of included data [38]. The resulting datasets usually excluded one or more of the 325 cells comprising the combined control (CON) and PMA- and LPA-treated (EXP) samples. Another procedure was to eliminate cells exceeding these limits based on the complete data represented in each trial. The first and second procedures were termed sample-by-sample or trial-by-trial removal, respectively.

Regularization is typically applied to all values of each descriptor represented in an experiment, but one can employ alternative datasets for mean-centering and scaling. The data of Trials 1-5 were regularized as described above. They were later combined into a single database totaling 1510 cells, which was used to autoscale the native descriptors’ values again. A third dataset, totaling 448 cells, was generated from a smaller number of cells treated with the solvent vehicle alone. This was made up from the Trials 1-5 CON samples together with other experimental controls generated by the same protocol [30, 39]. A fourth dataset, comprising 2623 cells, was made up from all experiments processed by the same protocol, some of which did not include a sample treated with PMA and LPA. A fifth dataset was accumulated from all cells processed by the high-resolution protocol, following transfection with exogenous plasmids [19, 31]. These accumulated samples provided a dataset of 624 cells. In all cases, autoscaling and subsequent computation of factor 4 values were performed as above.

## 3. Results

### 3.1 Values of the factor 4 trait following conventional mean-centering and scaling (autoscaling)

The process of autoscaling partially compensated for the heavy right tail of the values while centering the output around the mean of 0.0 (Supplementary Material, Figure S3). As expected, the EXP and CON samples within an experiment were indistinguishable (Figure 2B (left)). Five differences were found among repeated controls, due to the difference between the Trial 3 CON and all the others (Figure 2B (right)). Outlier removal on a sample-by-sample basis introduced a spurious difference between the CON and EXP samples in Trial 1 (Figure 2D (left)). Changing the basis of outlier removal to a trial-by-trial basis rendered the CON and EXP samples statistically indistinguishable again (Figure 2F (left)). Most differences among repeat CON samples remained when outliers were eliminated on either a sample-by-sample or trial-by-trial basis, as shown in Figure 2 (2B, 2D, and 2F (right)). Whereas removal of outliers overall eliminated <5% of total cells within a trial, sample-by-sample or trial-by-trial removal still eliminated 16.7% and 15.2%, respectively, of cells from a single sample (see Supplementary Material, Table S1).

**Figure 2.**
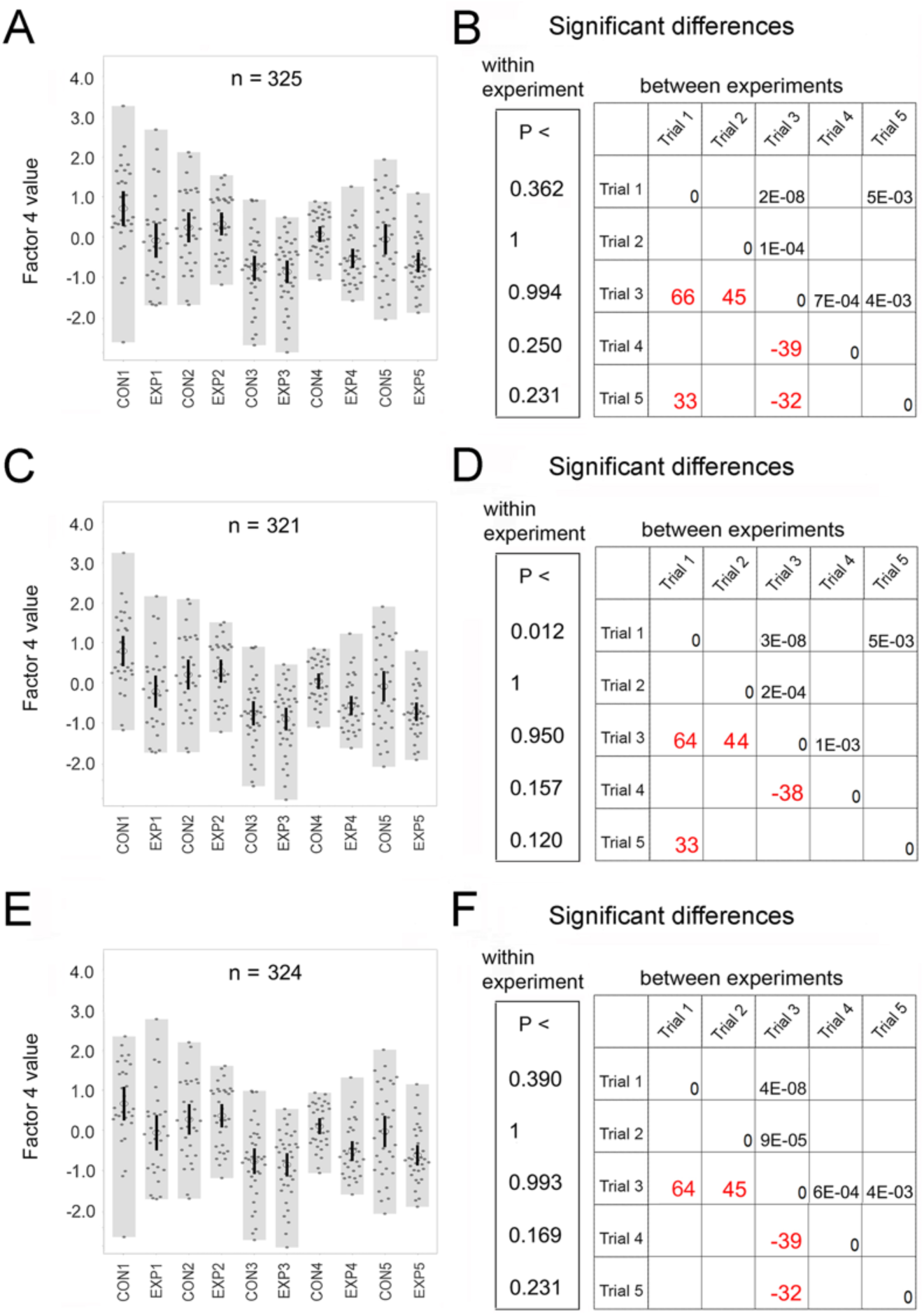
Factor 4 values after mean-centering and scaling with data from individual trials. (A, C, E) Values are shown as dots. Means are represented by a circle and 95% CIs by a vertical bar. (B, D, F) (Left) Significance of differences between the control and treated samples. (Right) Kruskal-Wallace test for differences among control samples in repeated trials shown by P-value in the upper right quadrant and mean rank score in red typeface. (A) Values without outlier removal. (B) (Left) P-values for control versus treated samples. (Right) Differences among CONs in the repeated trials. The number of cells per sample is: CON1=30, EXP1=30, CON2=30, EXP2=31, CON3=36, EXP3=36, CON4=32, EXP4=32, CON5=34, EXP5=34. (C) Values after outlier removal on a sample-by-sample basis. (D) (Left) P-values for CON versus EXP samples. (Right) Differences among CONs in repeated trials. One cell each is removed from samples CON1, EXP1, CON3, and EXP5. (E) Values after outlier removal on a trial-by-trial basis. (F) (Left) P-values for control versus treated samples. (Right) Differences among CONs in repeated trials. One cell is eliminated from the sample, CON1.

### 3.2 Contributions of asymmetric distribution and mean-centering to non-repeatability

The differences among repeated CON samples suggested that autoscaling the descriptors’ values within each trial may contribute to Type I errors. The true extent of repeatability is related to the important problem of batch effects, but an alternative explanation for non-repeatability might lie in the statistics of small samples. According to the Central Limit Theorem, if numerous trials were analyzed, the distribution of the CON means would be normal (Figure 3A). For any specific trial, however, its CON mean could fall anywhere within the normal distribution. This is due to the inherent variability of natural phenomena and is uncorrectable. Additional variability arose from the skewed distribution of factor 4 values. Factor 4 measures the prevalence of filopodia, and although the number displayed on a cell can vary, it cannot fall short of zero. Moreover, filopodia are formed by physiological processes, so factor 4 can rise to large values when cells overproduce the protrusions. Thus, the distribution is inherently asymmetric. When groups were made up from CON and EXP samples separately, both distributions were skewed (Figure 3B) showing that both contained cells overproducing filopodia. Certain other drug treatments caused the distribution to retract toward zero or stretch out into a larger range of values (Figure 3B (inset)). Large contributions to the heavy tail will increase the standard deviation used in autoscaling and decrease the variance within groups with more uniform distributions. It is likely the controls are among these groups, so any narrowing of their distributions could favor statistically significant distinctions between CON sample repeats.

**Figure 3.**
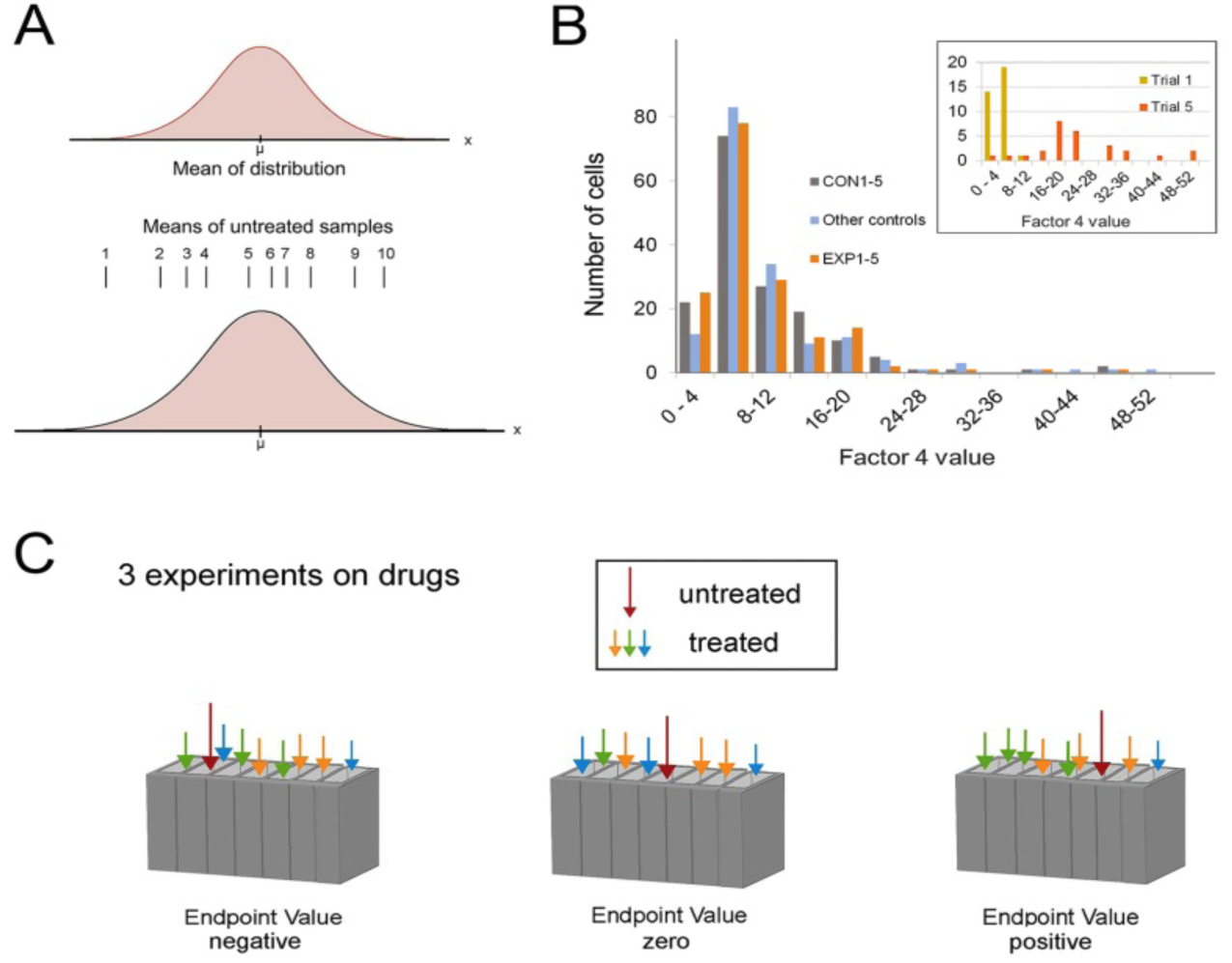
Effect of autoscaling. (A) Control means from repeated trials occupy a standard normal distribution (top). The trial means (1-10) are distributed in this range according to the probability function. (B) Distribution of values from pooled control and treated samples. High values occur in CON and EXP conditions in similar proportions. (Inset) Distribution of samples with the highest and lowest means in Trials 1-5. (C) The mean value of a scaled control takes a position relative to the remaining samples in that trial. Except for red, the colors represent different experimental reagents.

Mean-centering was another predictable source of variability. In addition to the EXP samples, the trials tested many other chemicals. The CON means would be biased to a low value if the samples had a high value on average. Conversely, if values were low overall, the CON mean would be biased to a high value (Figure 3C). Whereas ideally tests would expose cells to the same chemical entities under the same conditions [35], Trials 1-5 were from realistic experiments and did not test the same panel of drugs. Because mean-centering made all values within a trial interdependent, it could create a spurious difference among similar samples in repeated trials [40] and artifactually increase the difference between CON repeats. As investigators would rarely replicate the same set of treatments more than twice, this defect will generally be irremediable. In combination with the above trial-by-trial differences, this effect could exaggerate the differences between similar CON samples.

### 3.3 Effect on repeated controls (CON) of autoscaling with comprehensive databases

<u>In vitro</u> assay workflows automatically regularize the descriptors’ values, because it is assumed that it doesn’t change the relationships among samples in an experiment [41]. The results are considered an improvement, as the data are more normally distributed and thus, more tractable from a mathematical, statistical, and computational point of view [42]. For data regularized within trials, the mean of the Trial 3 CON sample was less than that of the other trials (Figure 2A, B). To the extent that the distinctions among CON samples depended on different distributions (see 3.2 Contributions of asymmetric distribution and mean-centering to non-repeatability), they would be erased by using a database with a greater range of values. This was tested by reprocessing the data from Trials 1-5 after replacing the individual trials’ databases with a pooled dataset of descriptors’ values. As the variance of the pooled database now represented all 1510 cells, the standard deviation used for autoscaling represented the full range for Trials 1-5. This had a marked effect on the CON sample of Trial 1. Its values had ranged from ∼3.3 to −2.7 (Figure 2A) but now only occupied half that range (Figure 4A). The differences between the Trial 3 CON sample and the other CON samples were no longer significant in data regularized with the 1510-cell database (cf. Figure 2B and 4B). This outcome was unaffected by outlier elimination (Figure 4D and F). Elimination of outliers on a sample-by-sample basis introduced the same spurious difference between the EXP and CON samples as found previously (cf. Figure 2D (left) and Figure 4D (left)).

**Figure 4.**
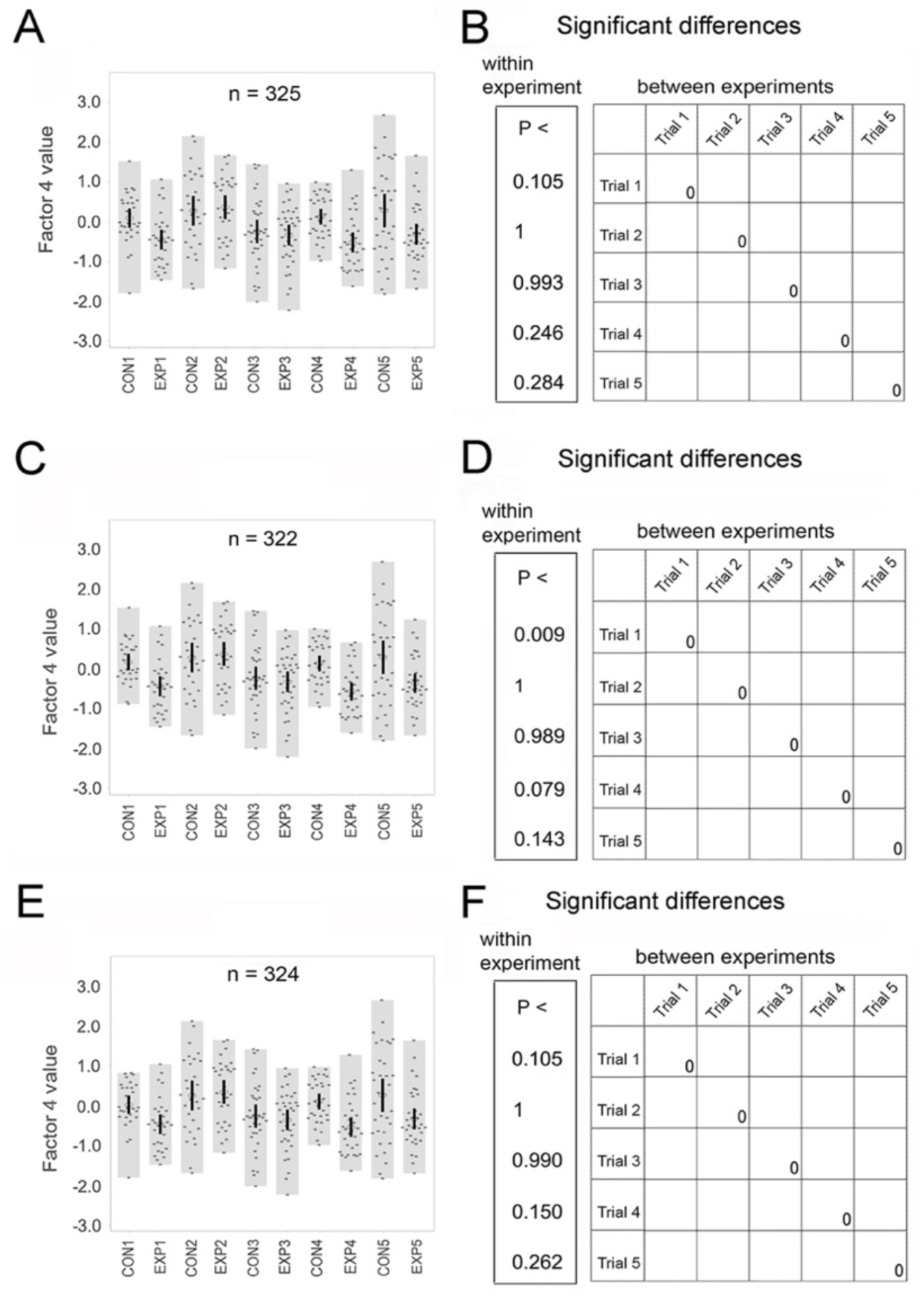
Factor 4 values after mean-centering and scaling with data from 1510 cells. (A, C, E) Values, means, and 95% CIs are shown as described in Figure 2. (B, D, F) (Left) Significance of differences between the control and treated samples. (Right) Kruskal-Wallace test for differences among CON samples in repeated trials shown by P-value in the upper right quadrant and mean rank score in red typeface. (A) Values without outlier removal. (B) (Left) P-values for CON versus EXP samples. (Right) Differences among CONs in the repeated trials. The number of cells per sample is given in the legend of Figure 2. (C) Values after outlier removal on a sample-by-sample basis. (D) (Left) P-values for control versus treated samples. (Right) Differences among CONs in repeated trials. One cell each is eliminated from sample CON1, EXP3, and EXP5. (E) Values after outlier removal on trial-by-trial basis. (F) (Left) P-values for the control versus the treated sample. (Right) Differences among CONs in repeated trials. One cell is eliminated from the sample, CON1.

A comparison of Figures 2 and 4 suggested that the difference between the Trial 3 CON mean and those of other trials was an artifact and a Type I error. The problem of autoscaling in HCAs was sometimes addressed by averaging the data from control wells. This was thought to reduce the impact of noise, edge effects, and similar variations [42, 43]. To determine whether restricting the database in this way would change the results, a new dataset was assembled from control samples alone (see 2.1 Data generation). The range of factor 4 values was affected very little by regularization to this dataset, totaling 448 cells, compared to the 1510-cell dataset (cf. Figures 4A and 5A). Moreover, the Trial 1-5 CON samples were still indistinguishable (Figure 5B (right), and the result was affected in a similar way by outlier elimination. Again, a spurious distinction between treated and control samples was introduced by outlier elimination on a sample-by-sample basis (cf. Figure 2D, 4D, and 5D (left)). The similarity suggested that the variance of the original descriptors’ values in the two datasets was similar. Indeed, for descriptors that weighed heavily in factor 4 computations, the standard deviations for the 448-cell dataset changed from only about 1% above to 19% below those for the 1510-cell dataset (data not shown). A larger dataset was also assembled, representing the descriptors’ values from all cells subjected to a similar experimental workup. All the comprehensive datasets, including the fourth set totaling 2623 cells, gave the same results (cf. Figures 4, 5, and Supplementary Material, Figure S4).

**Figure 5.**
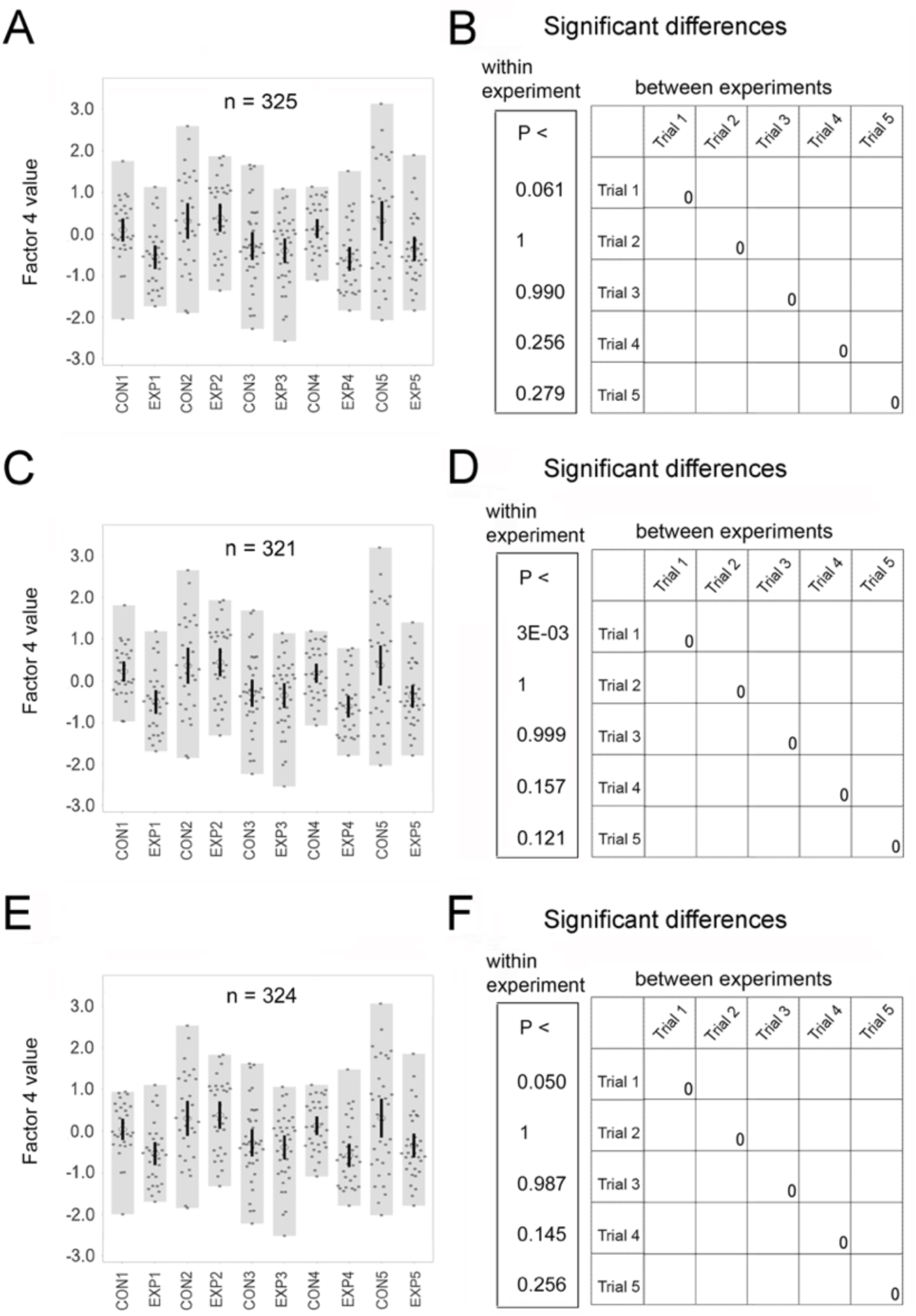
Factor 4 values after mean-centering and scaling with data from 448 control cells. (A, C, E) Values, means, and 95% CIs are shown as described in Figure 2. (B, D, F) (Left) Significance of differences between the control and treated samples. (Right) Kruskal-Wallace test for differences among CON samples in repeated trials shown by P-value in the upper right quadrant and mean rank score in red typeface. (A) Values without outlier removal. (B) (Left) P-values for CON versus EXP samples. (Right) Differences among CONs in the repeated trials. The number of cells per sample is given in the legend of Figure 2. (C) Values after outlier removal on a sample-by-sample basis. (D) (Left) P-values for control versus treated samples. (Right) Differences among CONs in repeated trials. One cell is eliminated from each of the samples, CON1, CON3, EXP4, and EXP5. (E) Values after outlier removal on experiment-by-experiment basis. (F) (Left) P-values for the control versus the treated sample. (Right) Differences among CONs in repeated trials. One cell is eliminated from the sample, CON1.

### 3.4 The effect of substituting different databases on classification patterns

<u>In vitro</u> assays are often designed to identify differences between a treated sample and the control(s). The classification of samples relative to one another within an experiment might be affected by the autoscaling procedure. As suggested above, a sample may reach the benchmark of P<0.05 if its distribution became compressed due to regularization with a comprehensive database. Indeed, regularization with the 1510-cell database yielded one new distinction over the results with individually regularized data (Figure 6 (red, top row)). Trials 2-4 showed no changes regardless of the database used for regularization (data not shown). Regularization to all comprehensive datasets gave identical outcomes (Figure 6).

**Figure 6.**
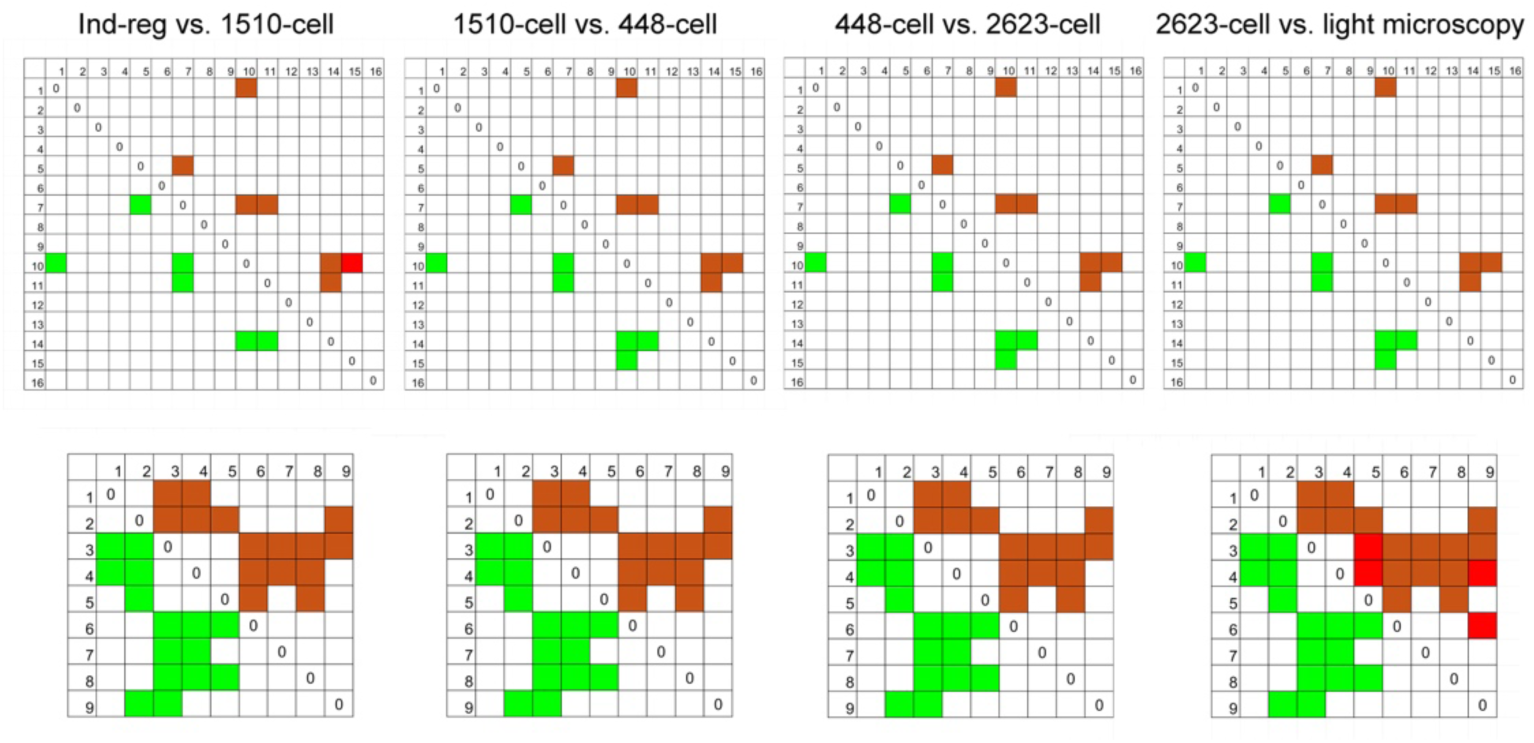
Classification patterns from autoscaling with substitute databases. New patterns are at top right along with any newly detected differences (red) over the old database (green, lower left). CON samples: Trial 1 (10) and Trial 5 (1). EXP samples are Trial 1 (2) and Trial 5 (2).

### 3.5 The effect of autoscaling to a database derived by a different procedure

It was important to compare the above patterns to those generated by autoscaling to the 800-cell database originally used to compute the scoring coefficients. The “light microscopy” dataset was generated from images of lower resolution, which were less precise than the images acquired in Trials 1-5. They were also derived from a more diverse group of cell lines. If the results diverged greatly from those described above, however, it would suggest that the model lacked stability. After autoscaling with the new database, one new difference appeared among repeated CONs compared to regularization with the other comprehensive databases (cf. Figures 4A-B, Figure 5A-B, and Figure 7A-B (right)). Outlier removal on a sample-by-sample basis introduced the spurious difference between Trial 1 CON and EXP samples noted previously (Figure 7D (left), while removal on a trial-by-trial basis restored both the above changes to the states observed before outlier elimination (Figure 7D and F (right)).

**Figure 7.**
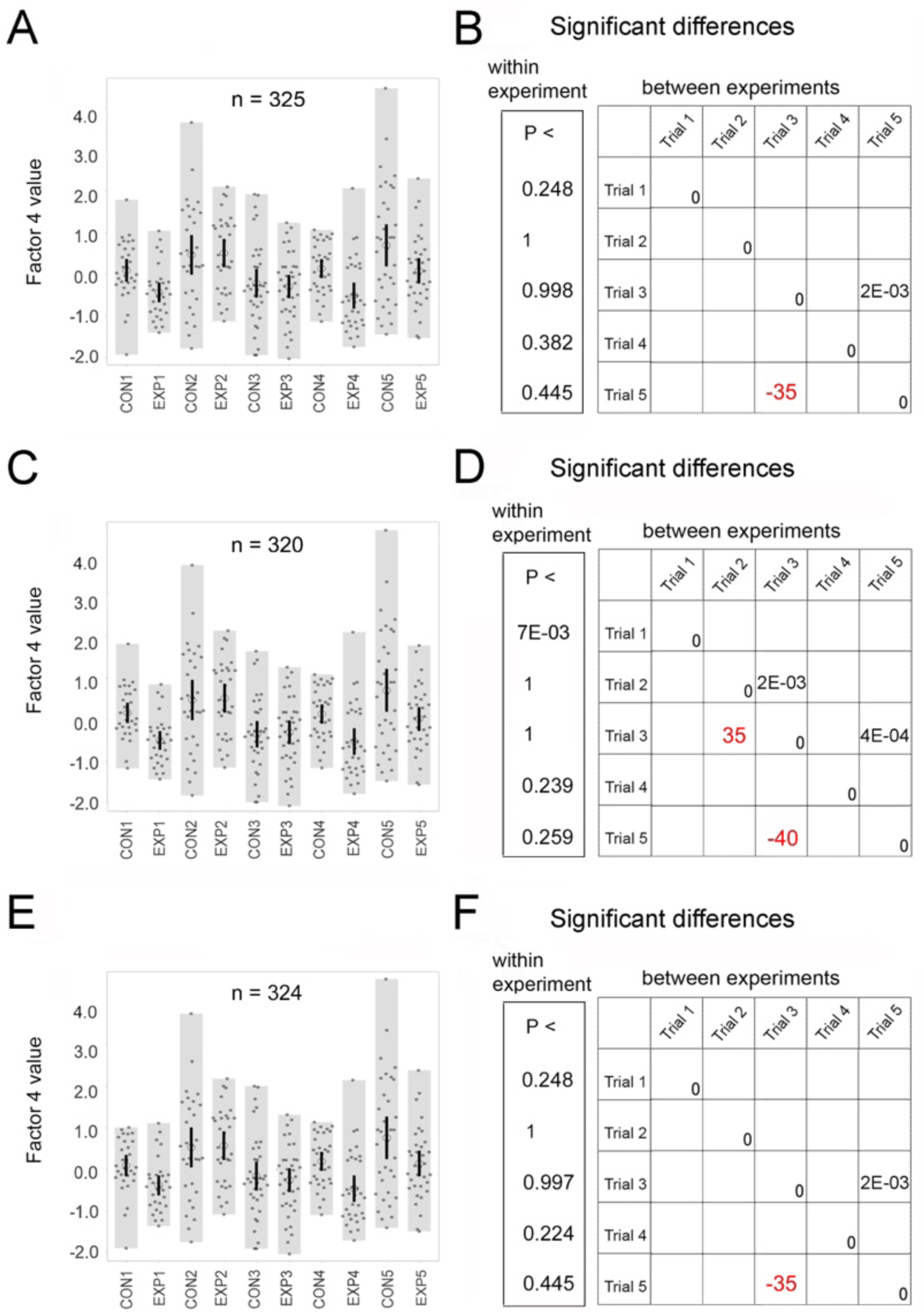
Factor 4 values after mean-centering and scaling with the light microscopy database. (A, C, E) Values, means, and 95% CIs are shown as described in Figure 2. (B, D, F) (Left) Significance of differences between the control and treated samples. (Right) Kruskal-Wallace test for differences among CON samples in repeated trials shown by P-value in the upper right quadrant and mean rank score in red typeface. (A) Values without outlier removal. (B) (Left) P-values for CON versus EXP samples. (Right) Differences among CONs in the repeated trials. The number of cells per sample is given in the legend of Figure 2. (C) Values after outlier removal on a sample-by-sample basis. (D) (Left) P-values for control versus treated samples. (Right) Differences among CONs in repeated trials. One cell is eliminated from each of the samples, CON1, EXP1, and EXP5. Two cells are removed from CON3. (E) Values after outlier removal on experiment-by-experiment basis. (F) (Left) P-values for the control versus the treated sample. (Right) Differences among CONs in repeated trials. One cell is eliminated from the sample, CON1.

When classification outcomes were analyzed, the only changes over regularization with the 1510-cell database occurred in Trial 5 (Figure 6A-F). Here, new differences occurred between samples 5 and 9 and four other samples. As none of the original outcomes were nullified, the results suggested that some differences merely rose to a new level of significance. Otherwise, the database substitution had no impact on classification. When a new database was composed from samples that had been transfected with exogenous plasmid vectors, the class assignments closely resembled those obtained by regularization with comprehensive datasets (Supplementary Material, Figure S5A-E). As in the case of regularization with the light microscopy dataset, the newly detected distinctions reflected differences that rose to a new level of significance.

### 3.6 The effect of scaling on repeatability of EXP sample means

Among the control samples, the Trial 3 CON had been distinguished from the others by its lower values (Figure 2A). The distinction was erased when a more comprehensive database was used for regularization (cf. Figures 2B, 4B, and 5B). If EXPs showed the same pattern, this would suggest it originated in environmental conditions peculiar to each experiment. The EXP samples showed a different pattern, however. The Trial 2 EXP mean had elevated values following regularization within individual trials. It differed significantly from the means of three other EXPs (Figure 8A), and the differences increased to four when the data were regularized to the 1510-cell database. Results after regularization to the 448- or 2623-cell datasets were the same (Figure 8B-E). As the overall pattern was dissimilar to that of the CONs, it could not be explained by the considerations of Figure 3A-C (see 3.2 Contributions of asymmetric distribution and mean-centering to non-repeatability). It was equally unlikely that the differences were due to conditions pertaining to any individual trial, because the CON and EXP means were statistically indistinguishable within each trial and were exposed to a similar environment within each trial. Thus, the different patterns must arise from other sources.

**Figure 8.**
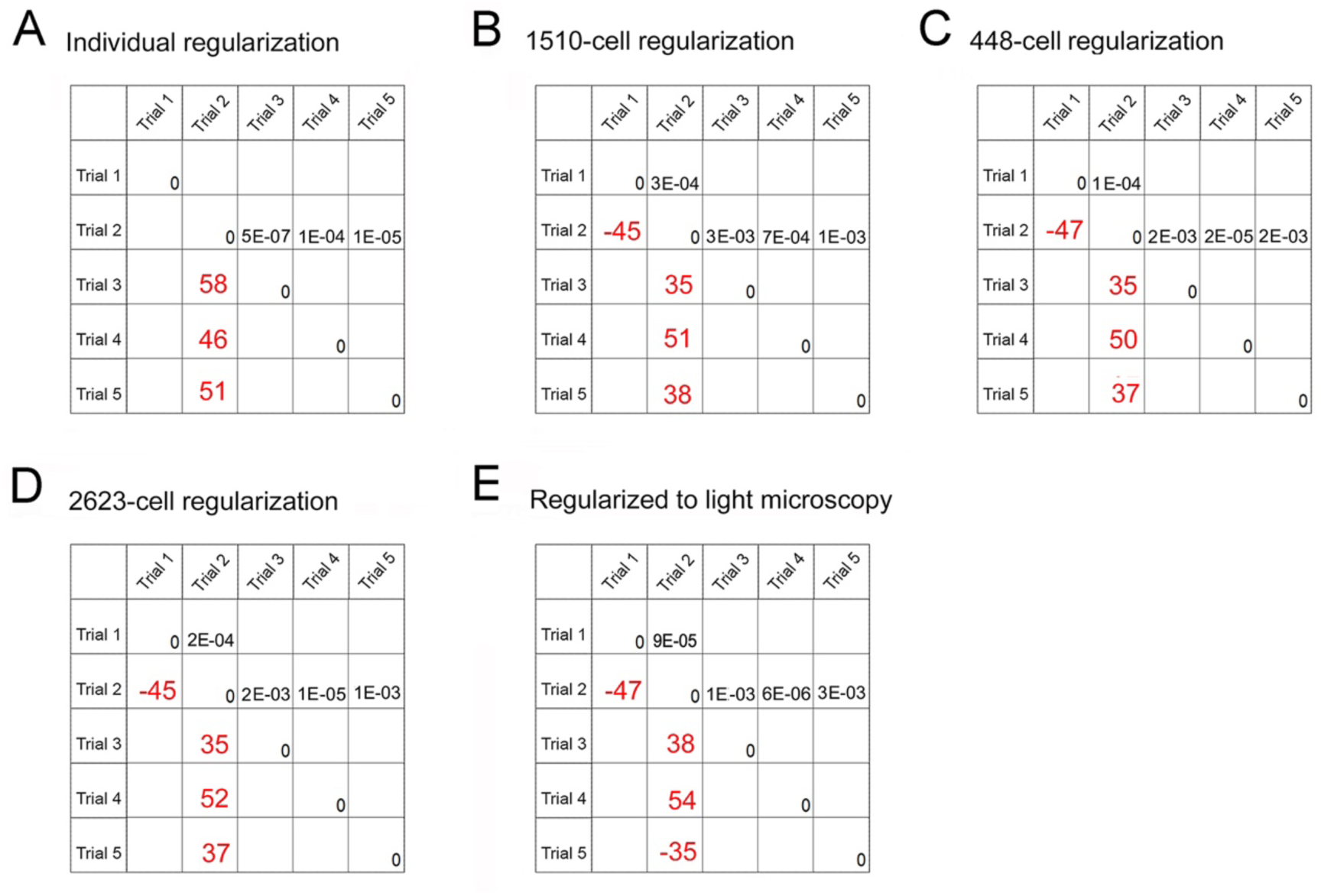
Differences among treated (EXP) samples after mean-centering and scaling with different databases. Results of the Kruskal-Wallace test are presented as described in the Figure 2 legend. (A) The Trial 2 EXP sample mean differs from those of Trials 3, 4, and 5 after autoscaling within each trial. (B-E) After autoscaling with the more comprehensive databases shown, the Trial 2 EXP differs from all other EXP samples.

When the pattern of classification of EXP samples was analyzed for Trial 1, it failed to follow the pattern of the CON (Figure S6A). In Trials 2-4, however, it exactly replicated the CON samples’ pattern (Supplementary Materials, Figure S6B-D). In Trial 5, two additional differences were detected for the treated (EXP) compared to the CON sample (Supplementary Materials Figure S6E). This suggested that the pipeline for data collection was reliable, and variations in the number and type of differences arose because differences that had been marginal rose to a new level of significance (see Discussion).

### 3.7 Outlier removal causes false negative (type II) and false positive (type I) effects

Outliers, i.e., entities whose values are out of the range of a preset statistic, are identified and removed in a typical HCA workflow. Various standards are used, but they may be applied on a sample-by-sample or a whole-population basis. Statisticians suggested that application at the sample level biases the data [42, 44]. In the current research, however, bias was introduced by both methods. When outliers were defined on a whole population basis, samples with a markedly right-shifted distribution were severely affected. Over 15% of a single sample could be removed (Supplementary Materials, Table S1) with the probable consequence of artifactually restricting range of values. This would cause false negative errors, i.e. missing a real difference between treated and control samples. The recommended practice, i.e., defining outliers relative to the entire population sampled, was not innocuous. The lost distinction between samples that really differed (Figure 9, cf. cyan squares at lower left and upper right) represented type II errors. Differences that had been recognized with all comprehensive databases were now missed. New distinctions between samples (Figure 9, red squares in upper right section) were false positives (Type I errors), as these differences were not recognized in data regularized to any of the three comprehensive databases (cf. Figures 6 and 9).

**Figure 9.**
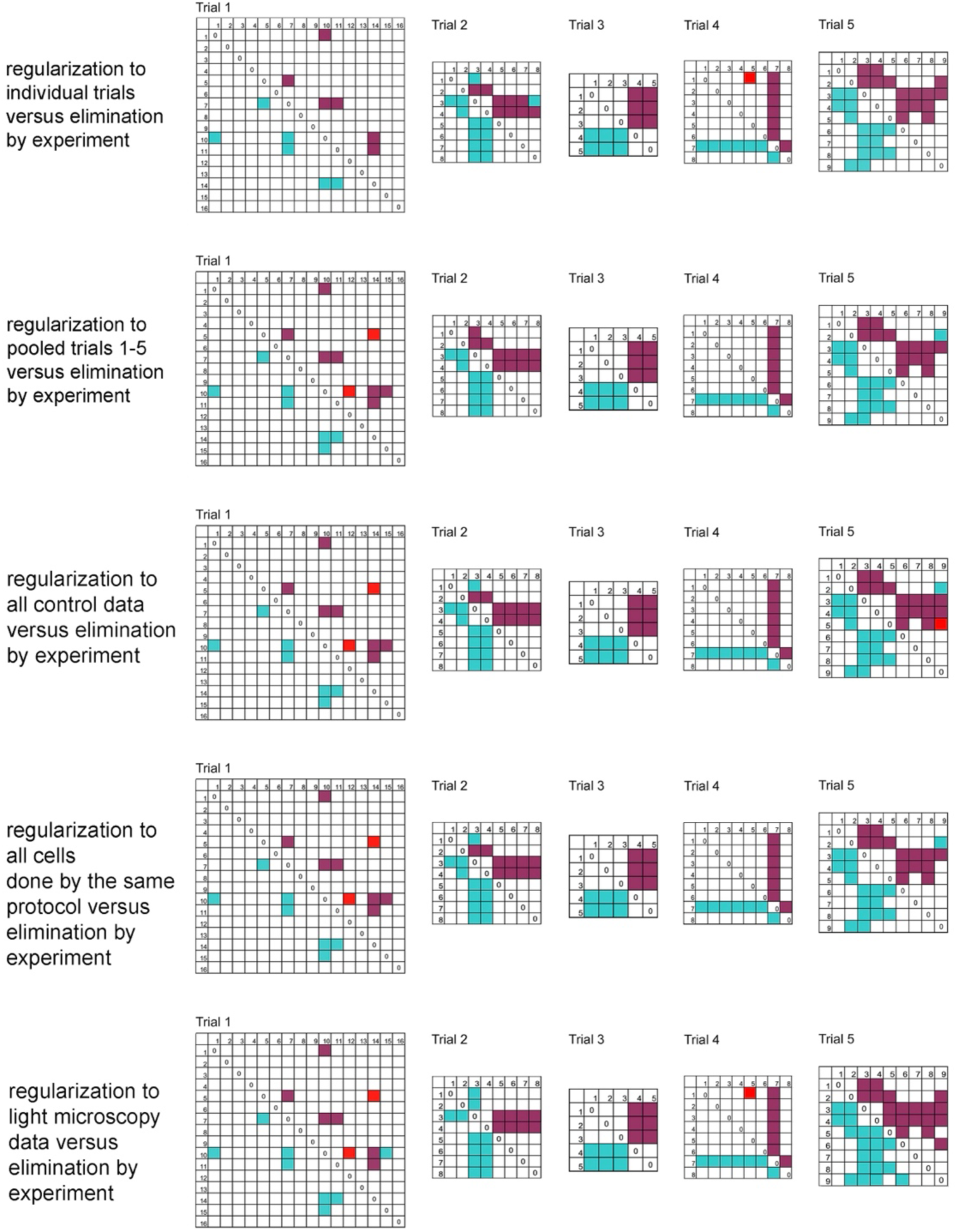
Effect on class assignments of outlier removal on a trial-by-trial basis. Significant differences detected after regularization to the datasets (cyan) are shown in the bottom left panels. They are regularized by individual trials, 1510-cell, 448-cell, 2623-cell, and the light microscopy datasets respectively. Significant differences found after outlier elimination are shown in the top right. Identical outcomes after outlier removal (magenta) are shown along with new differences created by outlier removal (red).

## 4 Discussion

### 4.1 Problems with image-based assay formats

Most assays used in drug discovery and development invoke algorithms to establish settings for each operation. Each one introduces a new set of vulnerabilities that can degrade the quality of the output, which are shown as hurdles in Figure 10. These processes present a problem for standardization and, more importantly, an obstacle to understanding cell phenotypes. Many methods have been proposed to deal with the complexity, but the gap in knowledge of how cell signaling networks, metabolism, and structure are intertwined is a barrier to effective investigation [8]. Workers investigating drug development expressed a need for phenotypes that are more relevant to diseases and for endpoints that are more tightly linked to phenotypes [9, 45]. The current results suggest that the effects of preprocessing on repeatability are subordinate to errors inherent in the image-based formats.

**Figure 10.**
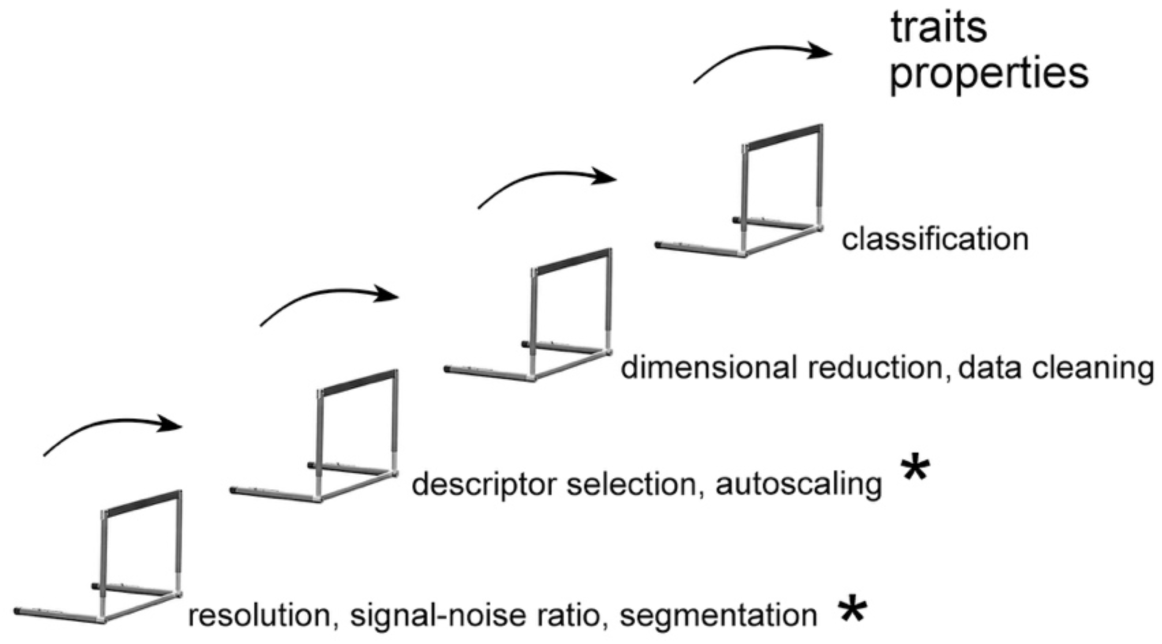
Areas of vulnerability in <u>in vitro</u> assays. Some (*) were minimized in this study: 1) labeling methods, 2) microscope settings, 3) segmentation, and 4) indeterminate endpoints. Factors common to all assay formats remain as the irreducible vulnerabilities here. These are unintended variations in compound concentrations, lot-to-lot variations in chemical purity, sample handling by different personnel, and non-repeatable conditions of the cell culture environment.

### 4.2 Effects of regularization (also called standardization, normalization, autoscaling, or Z-scaling)

As there is no reason to suppose that descriptors with high values are superior predictors, most microscopy-based assays include an autoscaling step. Its effect on reproducibility was studied by comparing the repeatability of CON and EXP sample means. After autoscaling within trials, the Trial 3 CON mean differed from the means of all other CON repeats. Among the EXP repeats, the Trial 2 mean exceeded those of most other EXP samples. Some instances of CON or EXP repeatability were affected by database substitution. However, the differences among the CON means were erased by substituting a more comprehensive database for regularization, which was not the case for repeated EXP samples. These differences couldn’t be rationalized by batch effects or the statistical factors shown in Figure 3, and their effect on classification outcomes was nil. For both CON and EXP samples, class assignments differed very little after autoscaling with different data. The results were similar when data from Trials 1-5 were regularized using: 1) combined control samples, 2) all samples from the same protocol, or 3) data acquired by a different protocol but processed similarly (Supplementary Material, Figures S5 and S6). Because the overall distributions of CON and EXP cells were similar (Figure 3B), the non-repeatable factors within a trial would affect descriptors alike in the CON and EXP samples.

As mentioned above, the current workflow controlled many of the environmental sources of variability, collectively called batch effects. The heterogeneity of variables acquired from the image were also minimized. The batch effects that were irreducible, together with small sample sizes and skewed distributions, may explain the non-repeatability of samples undergoing similar treatments.

### 4.3 Data cleaning

Descriptors’ values are measured directly from images in typical <u>in vitro</u> assays, but some are later eliminated as outliers (see for review [42, 46]). This may be innocuous when used to correct for variations in instrument performance, geometry of multi-well plates, etc. The ordinary rationale for eliminating descriptor values was that data in the tails were unrepresentative, but here, the descriptors’ values were considered data, so none were eliminated. Instead, values were eliminated from the low-dimensional datasets of factor 4 according to the most common, model-free method, the “Tukey fence.” Of the two variations of this procedure, the sample-by-sample elimination led to artificial (false positive) differences between the Trial 1 CON and EXP samples. The alternative was to define outliers by the whole range of values within an experiment. This led to both false positive and false negative differences in equal proportions (Figure 9). Because the range and distribution of factor 4 could stretch or shift on account of a drug treatment, a large fraction of cells with the highest or lowest values could be removed. Even with the most sparing method, nearly all the trials lost at least 3% of cells from a single sample (see Supplementary Material, Table S1). The fraction of cells eliminated varied by trial and by the method of defining outliers, but it was never small enough that the procedure could be considered harmless.

### 4.4 Classification

Although the methods of defining properties or traits by latent factor analysis are widely used in other fields, only a few groups have employed them in HCAs [6, 10, 47]. Because the derivation of loading factors for calculating their values favors the least noisy descriptors, the least robust ones tend to be screened out [17]. Classification by factor 4 values allowed sample-to-sample differences to be determined accurately and led to the finding that comprehensive databases generated by the same protocol all yielded identical patterns (see 3.3 Effect on repeated controls (CON) of autoscaling with comprehensive databases). The three other databases, namely those restricted to each trial, the 800-cell light microscopy database, or a database of transfected cells, yielded classifications only slightly different from the canonical pattern. This suggests that previous workers overemphasized formal repetition of sample statistics, as even an identical procedure, with an identical number of samples and treatments, will not give the same sample statistics for two repeats [48].

This study is a first approach to defining how each vulnerability affects the trait, factor 4, and contributes to the irreproducibility problem. It should be noted that a cell has many traits and properties, and the factor 4 trait is only a small portion of its phenotype. In our model, a combination of traits would define a single phenotype (Figure 1). Supplementary Figure S8 illustrates the concept of phenotype construction using three edge features of cells as an example. Other laboratories have defined 4-30 phenotypes based on images (Supplementary Material, Table S2). The number of theoretically possible classes can range up to 8E+32, which is larger than the number of stars in the known universe. Deriving low-dimensional variables may be required to allow accurate definition of traits and the subsequent derivation of phenotypes from traits.

Whereas a more accurate definition of traits may foster the development of image-based assays, it remains unclear the extent to which classification relies on having an identifiable, interpretable variable. Whereas it is not known whether other quantifiable properties resemble factor 4 in this respect, it seems likely. However, the variability of phenotypes cannot be foretold based on the current results.

## 5 Conclusions

Recent technical advances have made it possible to analyze images in ways that were never previously possible. Unfortunately, about half of reported experiments cannot be independently reproduced by another laboratory [49, 50]. It is likely that technical factors, called batch effects, contribute to this deficit, but there are few reports of real-world experiments where the technical factors are analyzed separately. The current work showed that variations in means and standard deviations measured in consecutive trials were subject to variations inherent in the assay conditions. However, the procedures ensured accurate class assignments despite these irreducible variations. This suggested that statistical classification was manageable, and the use of sample repeatability as a benchmark of assay quality is invalid. While it seems that quality and repeatability are unrelated, there is a lack of accepted criteria for judging the validity of results from image-based assays [49]. Although class assignments show promise for this purpose, if the controllable sources of variation summarized in Figure 10 were added back, e.g. signal/noise ratios, variations in instrument performance, and segmentation errors, the classification procedure may become unreliable. Further investigation of this aspect is warranted.

To summarize the main conclusions, they are:

1. Regularization can alter statistical significance in cases where small variations can affect the result, while outlier elimination consistently degrades data integrity.
2. Non-repeatable means can be attributed to irreducible technical factors (personnel, material purity and concentration, etc.), small sample sizes, skewed distributions, and possibly the way multidimensional data are reduced.

3 Class assignments are not affected by irreducible technical factors, such as materials, personnel, and non-repeatable environmental variables.

4 Classification pattern of an identifiable and interpretable trait is a sound criterion for assay quality.

Some practical recommendations about HCAs can also be made: 1) manage the controllable aspects of the assay format, 2) autoscale using any comprehensive database developed by the same protocol, 2) avoid outlier removal except for true artifacts, 3) use classification patterns instead of sample repeatability to evaluate assay quality.

## Abbreviations

ANOVA: analysis of variance
CON: control sample
EXP: sample treated with PMA and LPA
IQR: interquartile range
LPA: 1-oleoyl-sn-glycero-3-phosphate (lysophosphatidic acid)
P-value: probability of one group’s differing from another
PMA: phorbol 12-myristate 13-acetate
RNA: ribonucleic acid

## Declarations

### Clinical trial number

not applicable

### Ethics approval and consent to participate

Not applicable

### Consent for publication

Not applicable

### Availability of data and material

The 800-cell dataset is publicly available in ResearchGate. All datasets will .be made available to the editors and reviewers upon request and uploaded to a public site upon acceptance of the manuscript.

### Authors’ contributions

CH conceived the idea. CH analyzed the data, generated the figures, and wrote the manuscript.

## Acknowledgements

The author thanks her former students, Mr. Amarachintha, Mr. Demuth, Mr. Malwade, Mr. Roholt, Mr. Urban, Ms. Varghese, and Ms. Weber for generating the data used in the study. She also thanks Ms. Jessica Barnett and Ms. Kaitlyn Niek for editorial help.

A Novel Outlier Detection Algorithm Based on Symmetry and Distance Ratio

Zhai, HY; Fei, ZX and Ma, Y

27th International Conference on Pattern Recognition-ICPR-Annual 2025

PATTERN RECOGNITION, ICPR 2024, PT X

15310, pp.331-344

Outlier detection has become essential in various fields such as defense monitoring, fiscal anomaly identification, and business industries.

