## Supplementary_Materials for "Impact of Regularization Methods and Outlier Removal on Unsupervised Sample Classification"

| | | CON1 | EXP1 | CON2 | EXP2 | CON3 | EXP3 | CON4 | EXP4 | CON5 | EXP5 |
|---|---|---|---|---|---|---|---|---|---|---|---|
| Trial 1 | CON1 | 0 | | | | 1E-08 | 3E-09 | | 5E-04 | | 7E-08 |
| | EXP1 | | 0 | | | | | | | | |
| Trial 2 | CON2 | | | 0 | | 6E-05 | 2E-05 | | | | 2E-04 |
| | EXP2 | | | | 0 | 4E-06 | 1E-06 | | | | 1E-05 |
| Trial 3 | CON3 | 139 | | 98 | 112 | 0 | | 6E-05 | | | |
| | EXP3 | 145 | | 104 | 118 | | 0 | 8E-05 | | | |
| Trial 4 | CON4 | | | | | -89 | -95 | 0 | | | 6E-05 |
| | EXP4 | 80 | | | | | | | 0 | | |
| Trial 5 | CON5 | | | | | | | | | 0 | |
| | EXP5 | 133 | | 92 | 106 | | | | 83 | | 0 |

Supplementary Figure S1. Kruskal-Wallace test with Bonferroni corrections. The results of the post-hoc Dunn's test for differences is shown for five control and five treated sample distributions after autoscaling within individual trials. Significant differences are shown by P-value in the upper right quadrant and mean rank score in red typeface. The number of cells in each sample is CON1=30, EXP1=30, CON2=30, EXP2=31, CON3=36, EXP3=36, CON4=32, EXP4=32, CON5=34, EXP5=34. The control and treated samples within any single trial are indistinguishable on a statistical basis. These pairs are suitable for tests of errors, i.e., false positive and false negative results.

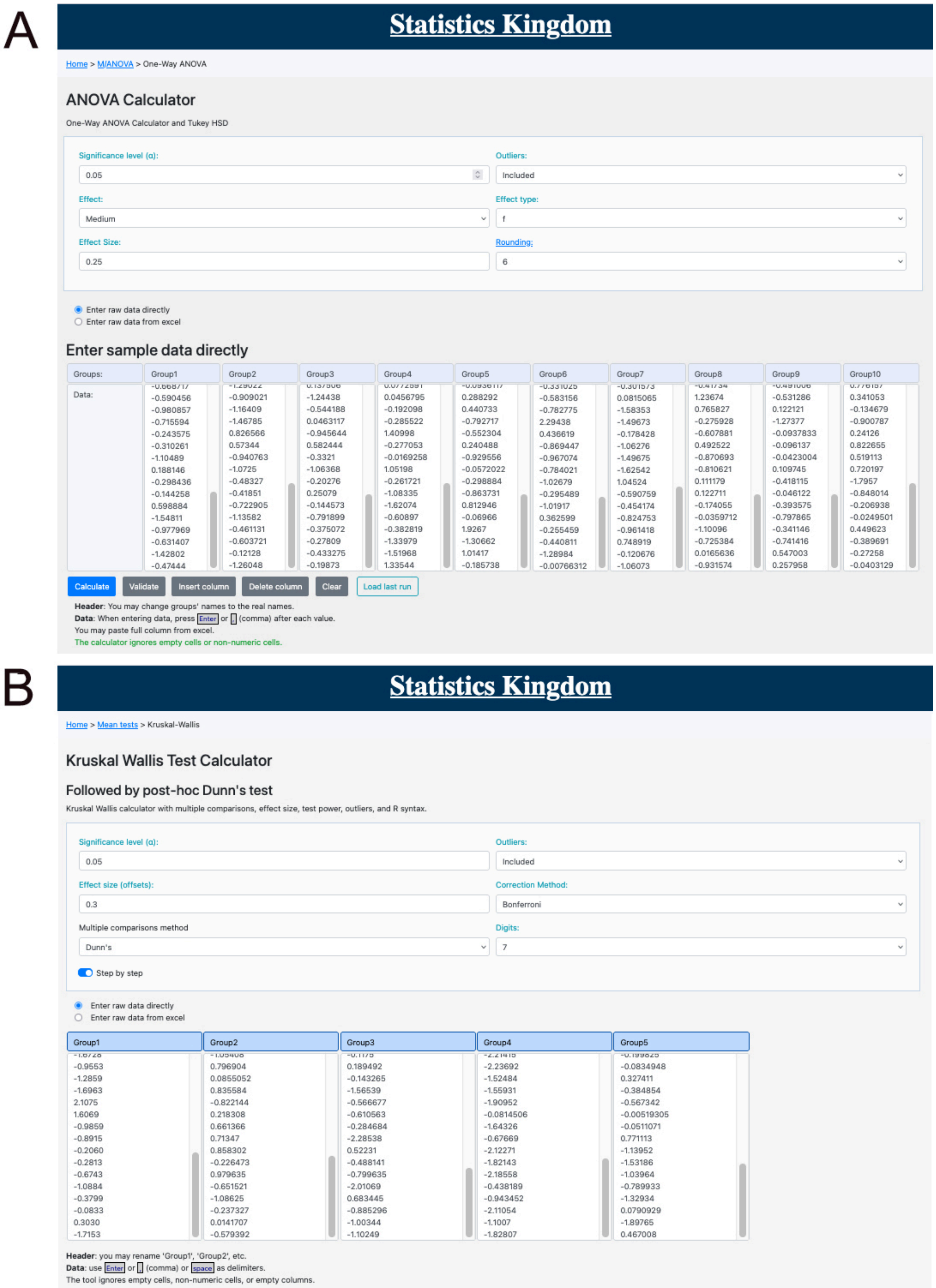


Figure S2. Online tests. (A) Settings for significance level (α), effect size (0.25), and effect type (f) in one-way ANOVA. (B) Settings for significance level (α), effect size, and correction for multiple comparisons in Kruskal-Wallis test. These can be reproduced in other programs for statistical analysis.

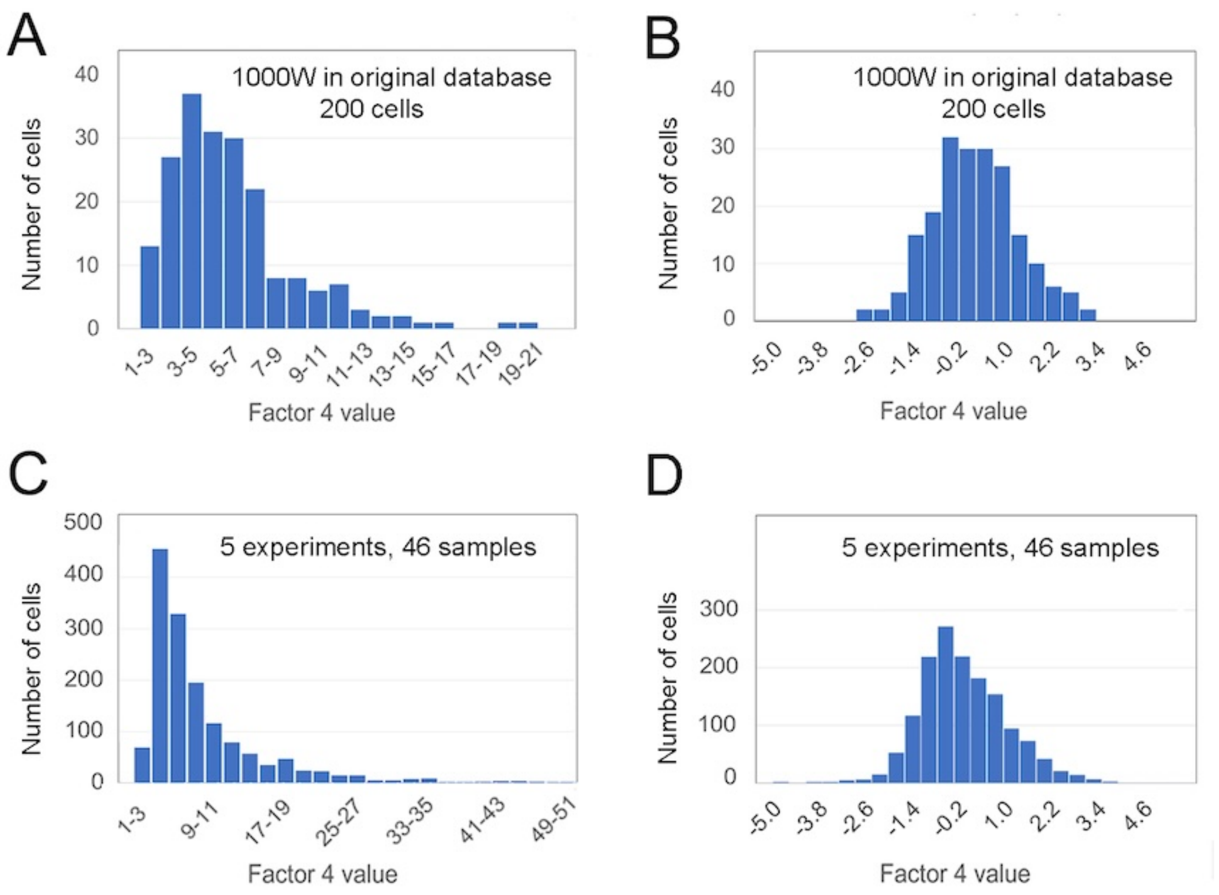


Supplementary Figure S3. Distribution of non-regularized and regularized factor values comparing image data from a low-resolution, high diversity database (A and B) with high-resolution, low-diversity images (C and D). (A) Non-regularized values of factor 4 for 1000W cells showing the long right tail of values. (B) Factor 4 values regularized by mean-centering and scaling within the 800-cell database. (C) Non-regularized values of factor 4 for the high-resolution, high-uniformity images of Trials 1-5 comprising 1510 cells. (D) Factor 4 values regularized by mean-centering and scaling with the pooled database of 1510 cells.

Table S1. Percentage of cells eliminated by outlier removal on sample-by-sample and trial-by-trial basis

| Database for regularization | Trial | Removal sample-by-sample | | Removal trial-by-trial | |
|---|---|---|---|---|---|
| | | Max %/sample | Average | Max %/sample | Average |
| Experiment by experiment | 1 | 16.7 | 4 | 6.1 | 2 |
| | 2 | 5.6 | 1 | 15.2 | 2 |
| | 3 | 5.6 | 2 | 5.6 | 1 |
| | 4 | 3.1 | 1 | 5.9 | 1 |
| | 5 | 3.7 | 1 | 0 | 0 |
| Average | 1-5 | | 1.7 | | 1.4 |
| Universal to pooled trials 1-5 | 1 | 6.1 | 2 | 6.7 | 2 |
| | 2 | 2.9 | 0 | 3.4 | 1 |
| | 3 | 8.3 | 2 | 5.6 | 1 |
| | 4 | 5.9 | 2 | 11.8 | 2 |
| | 5 | 3.7 | 1 | 3.2 | 0 |
| Average | 1-5 | | 1.6 | | 1.3 |
| Universal to controls (448 cells) | 1 | 13.3 | 3 | 6.7 | 3 |
| | 2 | 2.8 | 1 | 9.1 | 2 |
| | 3 | 8.3 | 2 | 5.6 | 1 |
| | 4 | 3.1 | 2 | 11.8 | 3 |
| | 5 | 6.4 | 2 | 3.2 | 0 |
| Average | 1-5 | | 1.9 | | 1.7 |
| Universal to all cells (2623 cells) | 1 | 16.7 | 3 | 6.2 | 3 |
| | 2 | 3.7 | 1 | 3.4 | 1 |
| | 3 | 2.8 | 1 | 5.6 | 1 |
| | 4 | 3.1 | 2 | 3.1 | 1 |
| | 5 | 3.7 | 1 | 3.2 | 0 |
| | 1-5 | | 1.5 | | 1.2 |
| Universal to original 800 cells | 1 | 16.7 | 3 | 6.7 | 3 |
| | 2 | 2.9 | 0 | 6.9 | 1 |
| | 3 | 8.3 | 3 | 8.3 | 2 |
| | 4 | 5.7 | 3 | 11.8 | 3 |
| | 5 | 3.7 | 1 | 0 | 0 |
| Average | 1-5 | | 2.2 | | 1.8 |

Cells removed from each of the samples and each of five trails were analyzed. Shown are the maximum percentage removed from any one sample and mean percentage removed from each trial.

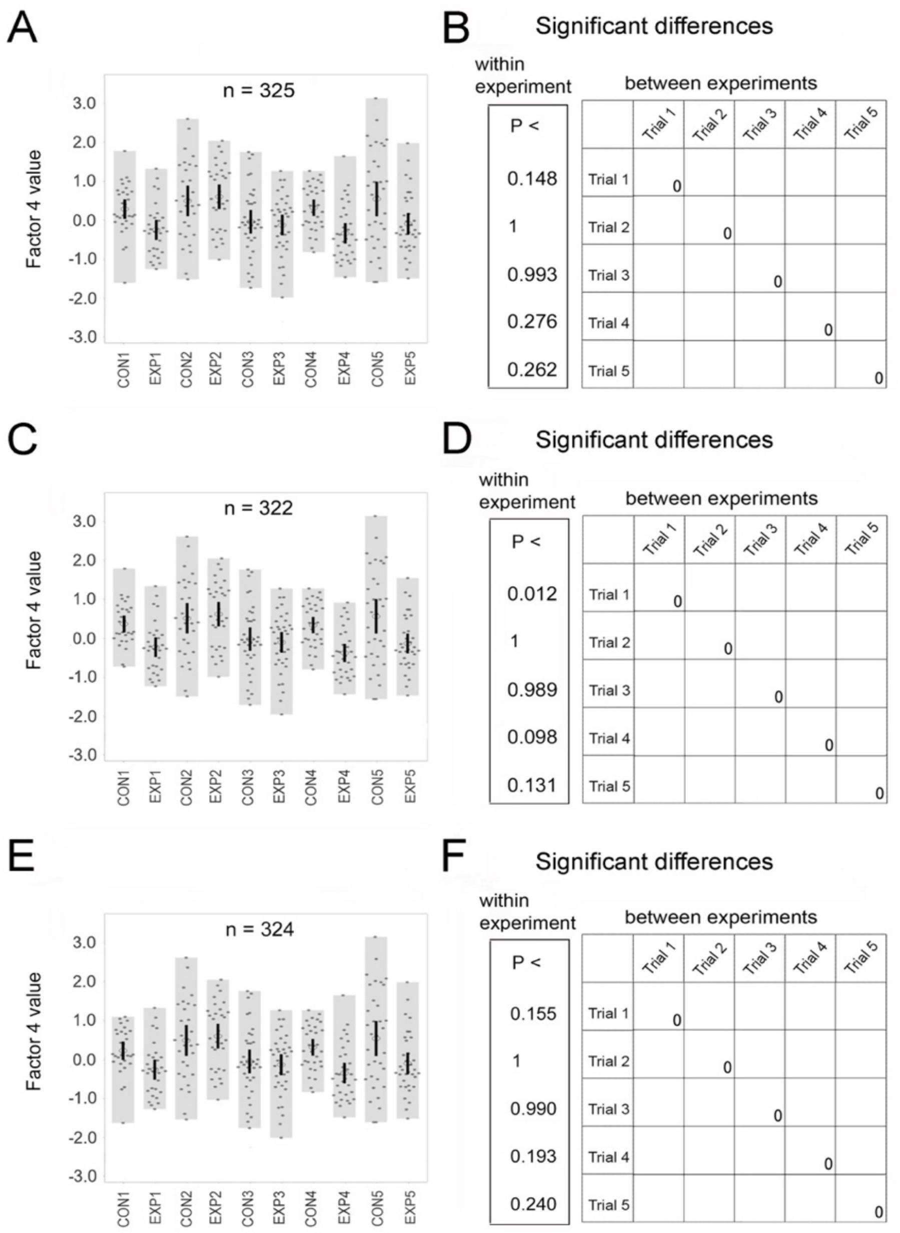


Supplementary Figure S4. Factor 4 values after mean-centering and scaling with data from 2623 cells. (A, C, E) Values, means, and 95% CIs are shown as described in Figure 2. (B, D, F) (Left) Significance of differences between the control and treated samples. (Right) Kruskal-Wallace test for differences among control samples in repeated trials. (A) Values without outlier removal. (B) (Left) P-values for control versus treated samples. (Right) Differences among controls in the repeated trials. The number of cells per sample is given in the legend of Figure 2. (C) Values after outlier removal on a sample-by-sample basis. (D) (Left) P-values for control versus treated samples. (Right) Differences among controls in repeated trials. One cell is eliminated from each of the samples, CON1, EXP4, and EXP5. (E) Values after outlier removal on trial-by-trial basis. (F) (Left) P-values for the control versus the treated sample. (Right) Differences among controls in repeated trials. One cell is eliminated from the sample, CON1.

Supplementary Figure S5. Classification outcomes from one-way ANOVA.  Factor 4 values after mean-centering and scaling with data from 624 cells transfected with exogenous DNAs (see Materials and Methods). Significant differences detectable after regularization with a comprehensive database (green) are shown in the lower left.  Identical outcomes after autoscaling with the transfection database are shown in the top right (brown) along with newly detected differences (red). All previous distinctions are detectable, and two are newly detected (red).

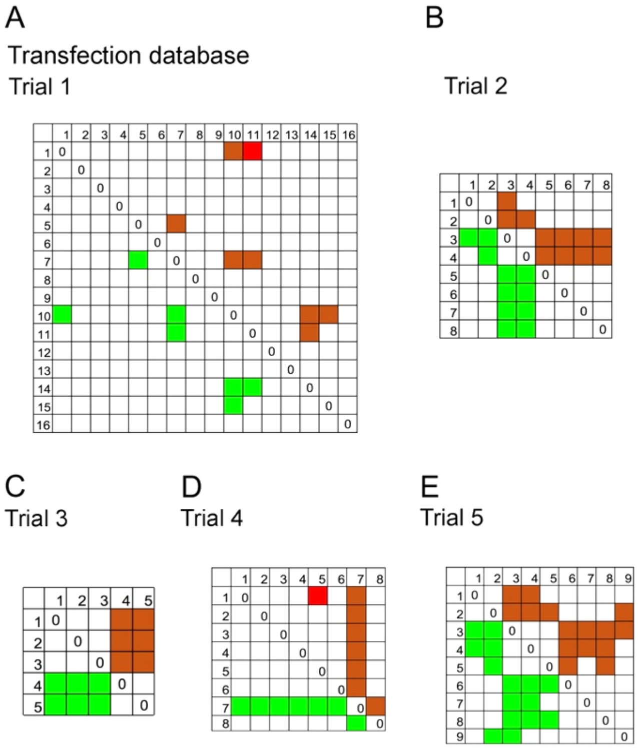


Supplementary Figure S6. Classification outcomes from one-way ANOVA. Differences detectable after regularization within each trial (green) are shown in the lower left.  Identical outcomes after autoscaling with other databases are shown in the top right (brown) along with newly detected differences (red). CONs are: Trial 1 (10), Trial 2 (8), Trial 3 (4), Trial 4 (8), Trial 5 (1). EXPs are: Trial 1 (2), Trial 2 (7), Trial 3 (5), and Trial 4 (1). (A) All previous distinctions are detectable, and one is newly detected. Trial 1 EXP (sample 2) is indistinguishable from all other samples. (B-D) All elements of the CONs pattern are reproduced by the EXPs. (E) Trial 5 EXP (sample 2) is significantly different from samples 5 and 9. (F) In Trial 5, four differences are newly detected after autoscaling with the light microscopy database.

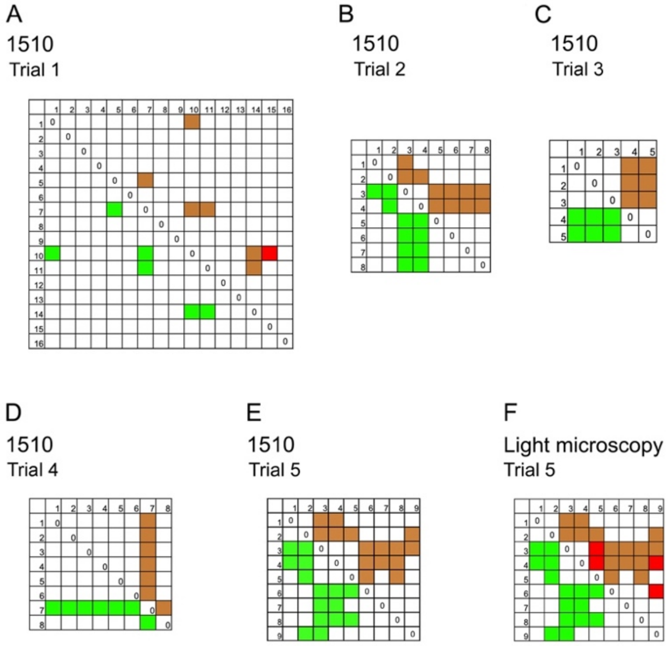

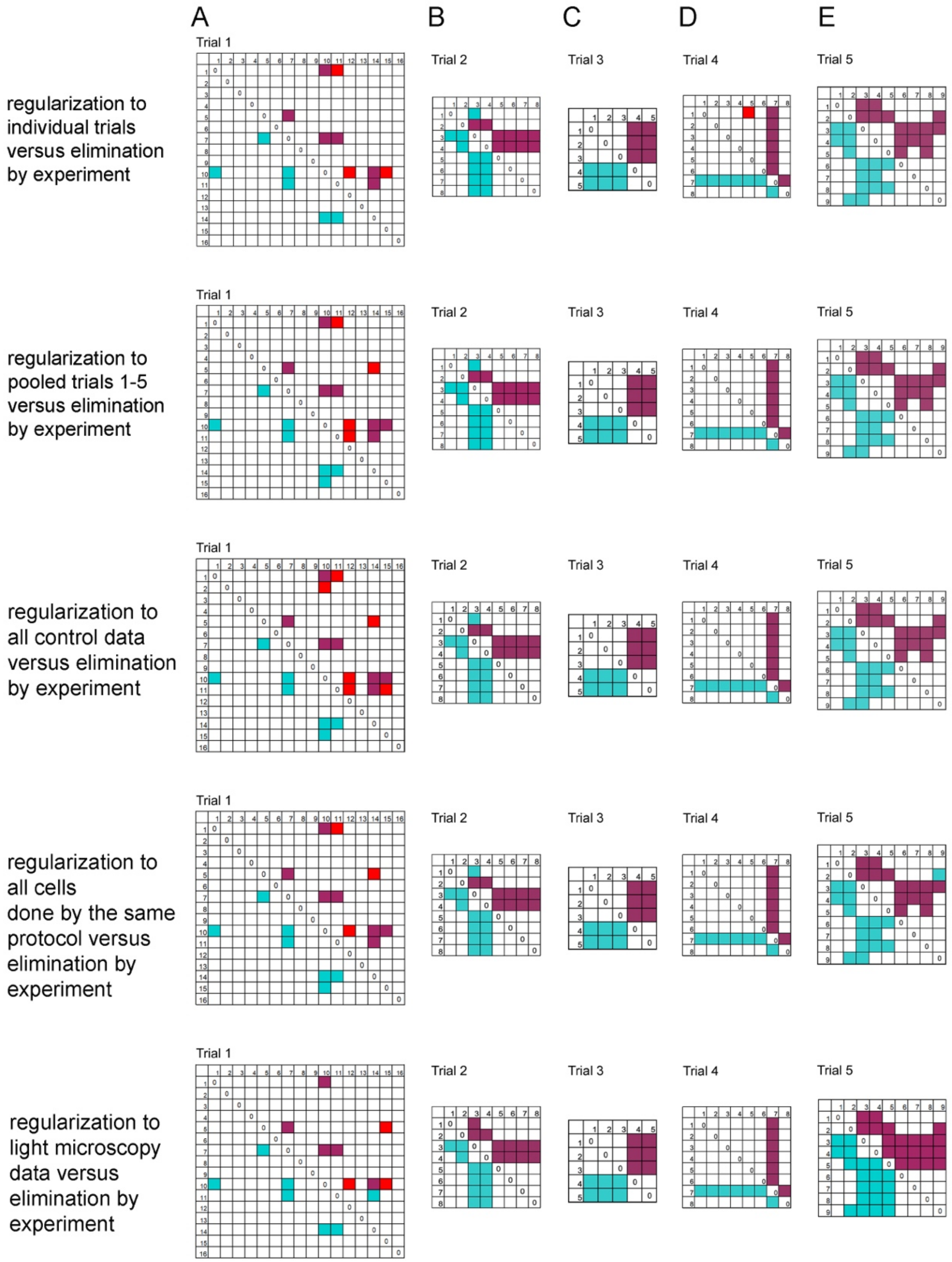


Supplementary Figure S7. Effect on class assignments of further definition of outliers by Z-scaling the autoscaled datasets. Significant differences detected after regularization to the datasets (cyan) are shown in the bottom left panels. They are regularized by individual trials, 1510-cell, 448-cell, 2623-cell, and the light microscopy datasets respectively. Significant differences found after outlier elimination are shown in the top right. Identical outcomes after outlier removal (magenta) are shown along with new differences created by outlier removal (red).)

Supplementary Figure S8. A cell phenotype can be portrayed by solving each trait of the treated sample by virtue of its factor value. Its mean can be significantly above, significantly below, or statistically indistinguishable from the control. The states are represented by green colors (above), orange-brown colors (below), or uncolored (indistinguishable), respectively. The traits are defined as features of the cell edge, filopodia, lamellipodia, and neurites respectively [22].

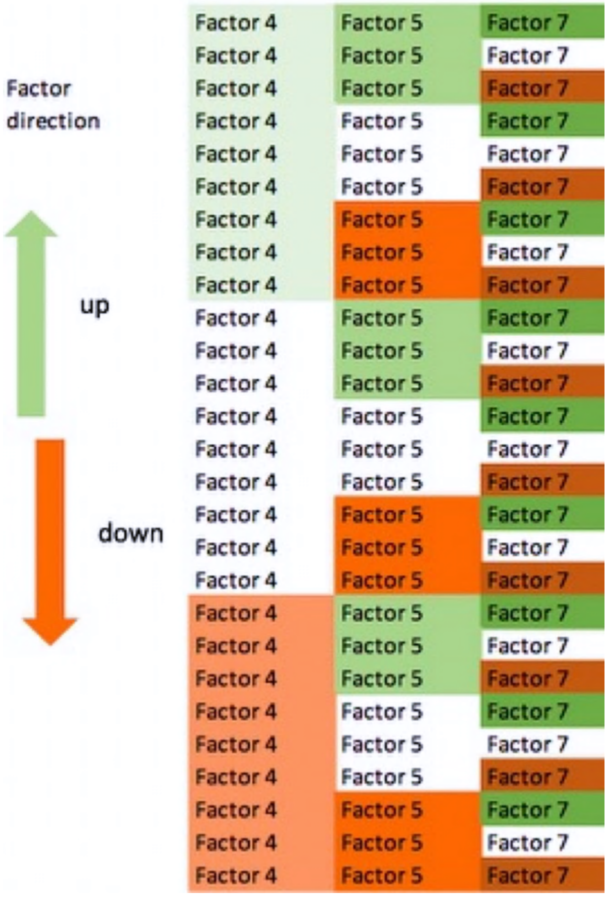


Table S2. Number of phenotypes or states by different classification methods

| Ref | Platform | # descriptors | # traits | # theoretical phenotypes | # number reported |
|---|---|---|---|---|---|
| [51] | Array Scan/Cellomics | 41 | 7 | 2187 | 4 |
| [52] | proprietary | 145 | 7[1] | 128 | 17 |
| [53] | Cellomics | 174 | 69 | 8.3E+32 | 4 |
| [54] | proprietary | 296 | 20 | 3.5E+09 | 30 |
| [55] | proprietary | 323 | 25 | 8.5E+11 | 11 |
| [47] | CellProfiler | 453 | 50 | 7.2E+23 | 12 |
| [56] | CellProfiler | 670 | 143 | 1.69E+68 | 18 |

The number of theoretical phenotypes is the number of combinations of traits or properties. Each trait exists in one of three possible states: significantly less in value, significantly more in value, or statistically indistinguishable from control.

[1]only presence or absence of the trait was possible, making the total number of classes 2^7.

References cited only in Supplementary Materials
